# Dysregulation of γδ intraepithelial lymphocytes precedes Crohn’s disease-like ileitis

**DOI:** 10.1101/2023.08.31.555710

**Authors:** Weili Xu, Natasha B. Golovchenko, Andrew Fong, Sajan Achi, Edie B. Bucar, Kaylynn J. Vidmar, Lanjing Zhang, Mark R. Frey, Immo Prinz, Theresa T. Pizarro, George Kollias, Karen L. Edelblum

**Author notes:** These authors contributed equally (Weili Xu, Natasha B. Golovchenko).

## Abstract

Intraepithelial lymphocytes expressing the γδ T cell receptor (γδ IELs) provide immunosurveillance of the intestinal barrier. Interestingly, γδ IEL number is reduced in patients with active Crohn’s disease (CD). Here, we report an underappreciated role for γδ IELs in maintaining mucosal tolerance during the onset and progression of CD-like ileitis using the TNF^ΔARE/+^ mouse model. Decreased epithelial HNF4G/BTNL expression is followed by a loss of ileal γδ IELs and impaired barrier surveillance prior to the histological onset of disease. A reduction of immunoregulatory CD39^+^ γδ IELs coincides with the influx of immature, peripheral CD39^-^ γδ T cells into the epithelium leading to an expansion of induced IELs, while an earlier depletion of γδ IELs correlates with accelerated onset of ileal inflammation. Our findings identify multiple layers of γδ IEL dysregulation prior to ileitis development indicating that the loss of tissue-resident immunoregulatory γδ IELs may contribute to the initiation of ileal CD.

## Introduction

Crohn’s disease (CD) is a chronic relapsing and remitting disease that affects the entire gastrointestinal tract with highest occurrence in the ileum. Genetic susceptibility, microbial dysbiosis, and immune dysregulation all contribute to the etiology of CD, yet the initiating events prior to disease onset or relapse remain unknown. Tumor necrosis factor (TNF) is a pro-inflammatory cytokine linked to increased epithelial permeability and apoptotic cell shedding^1,2^, both of which precede disease development^3–5^. Anti-TNF is widely used as a therapeutic in CD; however, patient response to treatment is highly variable^6^. Although it is difficult to evaluate the activation of intestinal immunity in individuals with CD prior to disease onset, further understanding of how mucosal immunity is affected during the initiating events in CD pathogenesis may be beneficial in designing novel therapeutic strategies to prevent disease relapse.

In contrast to experimental colitis, there are relatively few available animal models of spontaneous chronic ileitis^7^. Mice containing a deletion in the 3’ AU-rich region of *Tnf* (TNF^ΔARE^) exhibit increased transcript stability resulting in enhanced cytokine production^8^. Previous studies demonstrate that the TNF^ΔARE^ mouse model faithfully reproduces ileal pathology associated human CD, with heterozygous mice developing spontaneous ileitis at 8 weeks of age that progresses to transmural inflammation at 16 weeks^8^. While the progression of CD-like ileitis in TNF^ΔARE/+^ mice is well-described, alterations in mucosal immunity prior to the histological onset of disease are understudied.

Tissue-resident intraepithelial lymphocytes (IEL) are highly responsive to intestinal infection, injury, and inflammation^9,10^. The IEL compartment can be divided into two main subsets: natural IELs (CD8αα^+^ TCRαβ^+^ or TCRγδ^+^), which can be activated in an MHC-independent manner, and induced IELs that are antigen-experienced (CD4^+^ TCRαβ^+^, CD8αβ^+^ TCRαβ^+^ or CD4^+^ CD8αα^+^). While CD8αβ^+^ IELs contribute to ileal pathology in TNF^ΔARE/+^ mice through enhanced cytotoxicity due to endoplasmic reticulum stress^11–13^, the contribution of natural IELs, and more specifically γδ IELs, to ileitis remains poorly understood. γδ IELs dynamically survey the basement membrane and migrate within the lateral intercellular space (LIS) between neighboring enterocytes to function as sentinels of the epithelial barrier^14,15^. Recent studies revealed that the frequency of γδ IELs is reduced in patients with ileal CD^16^ and ulcerative colitis (UC)^17^, suggesting a protective role for γδ IELs in IBD. Although global deletion of γδ T cells results in enhanced severity of disease in experimental colitis^18–22^, the contribution of γδ T cells to chronic ileitis is less clear.

Prior studies investigating the requirement of γδ IELs in CD-like ileitis in TNF^ΔARE/+^ mice have yielded conflicting results^17,18^. TNF^ΔARE/+^ mice with a germline deficiency in γδ T cells exhibited similar disease development compared to controls^11^; however, αβ IELs can functionally compensate for the complete developmental absence of γδ IELs^23^. Administration of anti-TCRγδ to TNF^ΔARE/+^ mice exacerbated ileal pathology at 14 weeks of age^24^, yet this antibody induces TCRγδ internalization and only inhibits their adaptive functions^25^. Thus, this approach does not account for the myriad of innate-like functions attributed to γδ IELs^14,20,21,26,27^.

Herein, we profiled the ileal IEL compartment of TNF^ΔARE/+^ mice starting from 4 weeks of age and observed reduced frequency and absolute number of γδ IELs prior to ileitis onset. This early depletion of γδ IELs could be partly attributed to reduced signaling through the epithelial HNF4G/BTNL axis that is essential for γδ IEL homeostasis^28,29^. Further, intravital imaging demonstrated defective epithelial surveillance of the remaining γδ IELs preceding the initiation of ileal inflammation. Shortly after weaning, we observed a loss of CD39^+^γδ IELs and a concomitant influx of CD103^+^ CD39^-^ γδ T cells into the epithelium from the periphery. These findings suggest that decreased γδ IEL immunoregulatory capacity may contribute to a loss of mucosal tolerance as was evidenced by an increase in induced IELs during disease onset. Moreover, we found that premature depletion of γδ IELs correlated with accelerated disease development. Our study identifies multiple layers of dysregulation within the γδ IEL compartment early in the pathogenesis of CD-like ileitis and demonstrates that γδ IELs maintain mucosal tolerance to prevent ileal inflammation.

## Results

### The number of ileal γδ IELs is reduced prior to the initiation of chronic ileitis

The onset of ileitis in TNF^ΔARE/+^ mice has been previously reported at 8 weeks^8^; however, detailed immunological and histopathological analyses in the weeks leading up to ileitis onset have not been described. Scoring of ileal pathology from TNF^ΔARE/+^ mice collected at regular 1-2 week intervals starting at 4 weeks of age corroborate prior studies reporting that severe inflammation is first visible at 8 weeks (Figure 1a,b). Histological scores were driven largely by the extent of immune cell infiltration, with increases in submucosal and transmural inflammation observed as disease progressed (Figure 1c).

**Figure 1.**
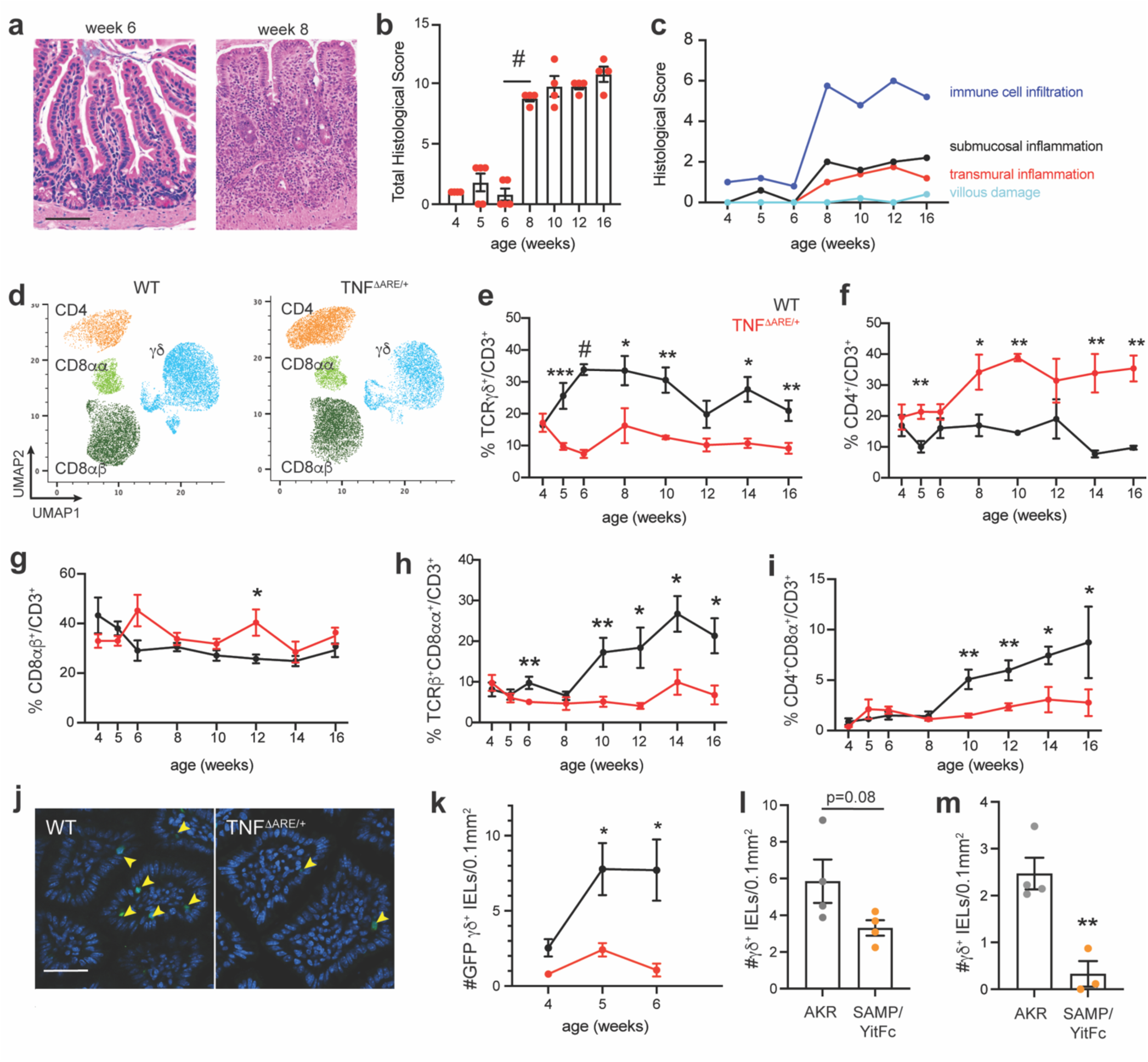
A reduction of ileal γδ lELs is observed prior to ileitis development in TNF^ΔARH+^ mice. (a) H&E micrographs of ilea from TNF^ΔARB+^ mice at 6 and 8 weeks of age. Scale bar = 100 pm. (b) Total and (c) individual components of histological scores of ilea from TNF^ΔARB+^ mice 4-16 weeks of age. (d) UMAP analysis of ileal lELs in 8-week-old WT and TNF^ΔARE/+^ mice. Frequency of ileal (e) TCRγδ+, (f) CD4+, (g) CD8αβ+, (h) CD8αα+ and (i) CD4+ CD8αα+ lELs (gated on CD3+) in WT and TNF^ΔARE/+^ mice from 4-16 weeks of age. (j) Immunofluorescence images of 5-week-old ilea from TcrdGDL (WT) and TcrdGDL; TNF^ΔARE/+^ mice. γδ T cells are shown in green and nuclei in blue. Scale bar = 30 µm. (k) Morphometric analysis of ileal GFP+ γδ lELs in WT and TNF^ΔARE/+^ mice from 4-6 weeks of age. *In situ* hybridization for *Trdc* in AKR and SAMP/YitFc mice at (I) 4 and (m) 10 weeks of age. n = 3-9. All data are shown as mean ± SEM from at least 2 independent experiments. Each data point represents an individual mouse. Statistical analysis: unpaired t-test. *P<0.05, *‘P<0.01, ***P< 0.001, #P< 0.0001.

To investigate whether IEL composition is altered preceding and during disease initiation, we profiled the IEL compartment of TNF^+/+^ (WT) and TNF^ΔARE/+^ mice over the same time course (Figure S1a). UMAP analysis shows a reduction in the frequency of γδ IELs with a concomitant increase of CD4^+^ TCRβ^+^ IELs at 8 weeks of age in TNF^ΔARE/+^ mice, indicating alterations in IEL subsets at the onset of ileal inflammation (Figure 1d). The earliest change within the IEL compartment in TNF^ΔARE/+^ mice was the pronounced reduction in frequency of γδ IELs at 5 weeks of age (Figure 1e). CD4^+^ IELs were substantially increased at the onset of inflammation (Figure 1f), followed shortly thereafter by an increase in induced CD8αβ^+^ IELs (Figure 1g, Figure S1b). The latter findings are in line with previous reports highlighting UPR-induced ER stress within CD8αβ^+^ TCRβ^+^ IELs promoting disease pathogenesis in the TNF^ΔARE/+^ model^13^. At 10 weeks, a reduction in natural CD8αα^+^ TCRβ^+^ (Figure 1h, Figure S1c) and regulatory CD4^+^ CD8αα^+^ IELs (Figure 1i) becomes apparent. These data highlight the profound shift in the IEL compartment from populations with regulatory and innate-like properties toward an antigen-specific adaptive response over the course of disease progression.

Since flow cytometry provides only the relative frequency of individual populations and there is variable efficacy in isolating IELs from the gut, we crossed TNF^ΔARE/+^ mice to those expressing a γδ T cell reporter and performed morphometric analysis to quantify ileal γδ IEL number. We found a stark reduction in the number of ileal γδ IELs in TNF^ΔARE/+^ mice compared to WT (Figure 1j,k). To determine whether this decrease in γδ IEL number is unique to this mouse model or a common feature of CD-like ileitis, we used SAMP1/YitFc (SAMP) mice^30,31^ and performed *in situ* hybridization to assess γδ IEL number prior to, and during the onset of, ileal inflammation. SAMP mice develop early epithelial alterations at 3-4 weeks of age, before histologic evidence of ileitis, whereas active inflammation is clearly evident by 10 weeks of age compared to age-matched AKR (parental control) mice^32^. We find that at 4 weeks of age, there is a slight reduction in γδ IEL number in SAMP mice compared to AKR littermates which is more pronounced at 10 weeks (Figure 1l,m). Taken together, our results show a specific reduction in γδ IEL frequency and number weeks before the onset of chronic inflammation in two different models of CD-like ileitis.

### Reduced signaling through the epithelial HNF4G/BTNL axis in TNF^ΔARE/+^ mice correlates with impaired γδ IEL homeostasis

Based on these findings, we posited that the loss of γδ IELs in TNF^ΔARE/+^mice may be attributed to reduced proliferation or enhanced cell death. Consistent with this, EdU incorporation was decreased and the frequency of apoptotic γδ IELs was increased in 5-week-old TNF^ΔARE/+^ mice relative to WT (Figure 2a-d). Epithelial butyrophilin proteins (BTNL) are critical for the maturation and development of γδ IELs^28^, and these genes were recently shown to be regulated by HNF4A and HNF4G transcription factors^29^. A reduction in *Hnf4g* was apparent at 4 weeks of age in the ilea of TNF^ΔARE/+^ mice, yet *Hnf4a* expression was not affected (Figure 2e). Subsequently, we observed decreased *Btnl1* and *Btnl6* at 5 weeks of age (Figure 2f). We next asked whether the distribution of *Btnl1/6* was altered along the crypt-villus axis at these early timepoints in TNF^ΔARE/+^ mice. Although *in situ* hybridization confirmed our qPCR results, no distinct regional differences in gene expression were noted (Figure 2g, Figure S2a,b). Together, these results indicate that reduced signaling through the HNF4G/BTNL axis adversely affects maintenance of the γδ IEL compartment resulting in reduced proliferation and increased apoptosis.

**Figure 2:**
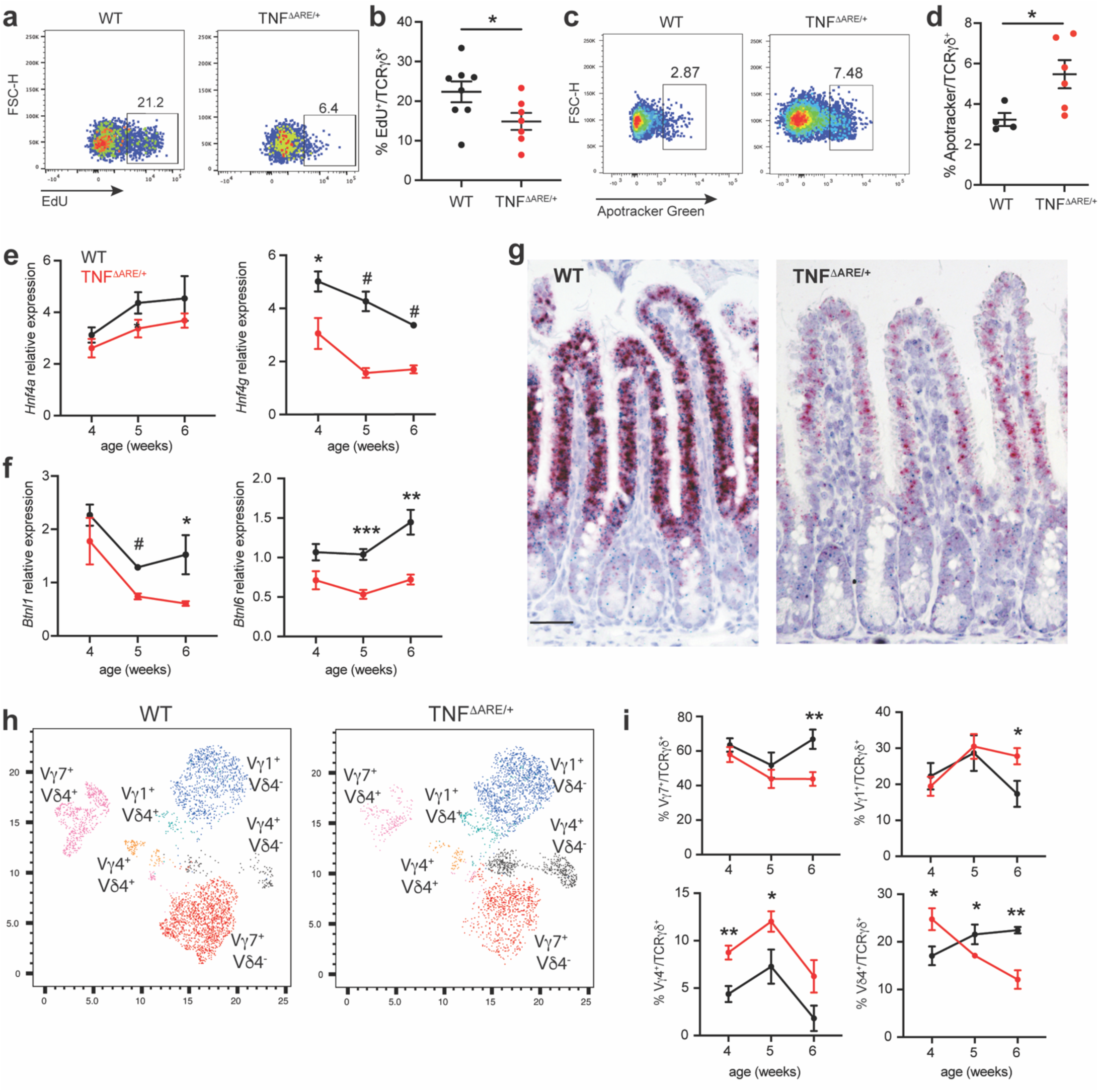
Early defects in the epithelial HNF4G/BTNL signaling axis in TNP^ΔRE/+^ mice negatively impact γδ IEL homeostasis. (a) Representative plot and (b) frequency of EdU+ γδ lELs isolated from the ilea of 5-week-old WT and TNF^ΔARE/+^ mice. (c) Representative plot and (d) frequency of apoptotic γδ lELs in 5-week old WT and TNF^ΔARE/+^ mice. Relative expression of ileal (e) *Hnf4a, Hnf4g,* (f) *Btnl1* and *Btnl6* normalized to *Cdh1.* (g) In situ hybridization showing *Btnl1* (green) and *Btnl6* (red) expression along the crypt-villus axis in 6-week-old WT and TNF^ΔARE/+^ mice. Scale bar =100 µm. (h) UMAP analysis and (i) frequency of Vγ and Vδ subsets among ileal lELs in WT and TNF^ΔARE/+^ mice from 4-6 weeks of age. n=4-8. All data are shown as mean ± SEM from at least 2 independent experiments. Each data point represents an individual mouse. Statistical analysis: unpaired t-test. *P<0.05, **P<0.01, ***P< 0.001, #P< 0.0001.

Loss of BTNL1/6 expression influences the γ and δ chain usage among γδ IELs, which are a heterogenous population^28^. Vγ7 is the predominant Vγ chain within the IEL compartment, followed by Vγ1 and Vγ4, with the majority of these IELs being Vδ4-negative. Therefore, we next assessed whether the proportion of γδ IEL subsets was altered in TNF^ΔARE/+^ mice in the weeks leading up to disease onset. At 6 weeks of age, there was a notable decrease in Vγ7^+^ and Vδ4^+^ IELs accompanied by an increase in Vγ1^+^ IELs in TNF^ΔARE/+^ mice relative to WT (Figure 2h,i). Interestingly, the frequency of Vγ4^+^ IELs was increased across all early time points in TNF^ΔARE/+^ mice leading us to hypothesize that there may be a selective reduction in cell proliferation or increased apoptosis among Vγ7^+^ γδ IELs. While no changes within the Vγ7^+^ population were observed in 6-week-old TNF^ΔARE/+^ mice, proliferation was impaired in Vγ1^+^ and Vγ4^+^ IELs and apoptosis enhanced in Vγ1^+^ IELs (Figure S2c,d). Collectively, these data indicate that impaired HNF4G/BTNL signaling in the ileum of TNF^ΔARE/+^ mice results in a shift within the IEL compartment toward Vγ7^-^ IELs; however, the increase in Vγ1^+^ and Vγ4^+^ IELs is not due to a local expansion within the epithelium.

### Impaired γδ IEL surveillance of the epithelial barrier precedes ileitis onset

Previous work from our group demonstrates that γδ IELs play a critical role in immunosurveillance in the small intestine^14,15,33^, thus we next asked whether the surveillance behavior of the remaining γδ IELs is affected prior to disease initiation. To this end, we performed intravital microscopy on the ileal mucosa of 6-week-old TcrdGDL (WT) and TcrdGDL;TNF^ΔARE/+^ mice. The reduction in γδ IEL number in TNF^ΔARE/+^ mice was again evident in comparison to age-matched WT littermates (Figure 3a). Although overall displacement was not impacted (Figure 3a), the mean track speed of γδ IELs was substantially reduced in TNF^ΔARE/+^ mice (Figure 3b). This reduced motility could not be attributed to an increase in cell arrest along a given track, yet we observed a trend toward increased dwell time of individual γδ IELs within the LIS in TNF^ΔARE/+^ mice (Figure 3c,d). Interestingly, γδ IEL migratory behavior was similar between 5-week-old littermates of either genotype (Supplementary Figure 3).

**Figure 3:**
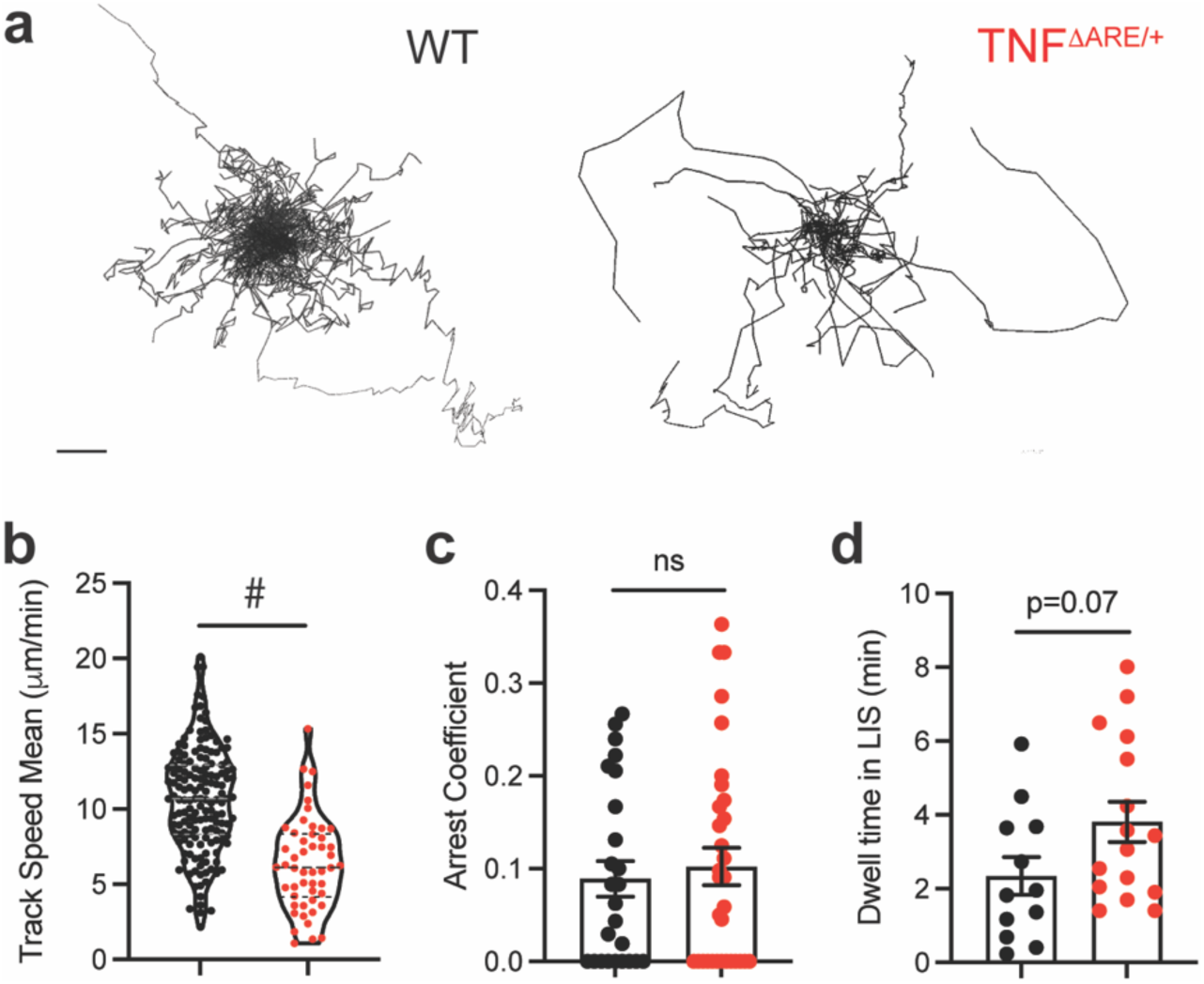
Reduced γδ IEL surveillance of the epithelial barrier precedes ileitis onset. Intravital microscopy was performed on ileal mucosa of 6-week-old TcrdGDL TNF^+/+^ (WT) or TcrdGDL;TNF^ΔARE/+^ mice. (a) Rose diagrams illustrating displacement of γδ IELs overtime. Scale bar = 10 µm. (b) Mean track speed, (c) arrest coefficient and (d) dwell time of individual γδ IELs within the LIS are shown. (b) Median and quartiles are shown (c,d) Data represent the mean ± SEM from 3 mice of each genotype. n= 155 tracks (WT), 45 tracks (TNF^ΔARE/+^). #P< 0.0001

Based on our previous studies showing that γδ IEL migration is dependent on expression of the tight junction protein occludin^14,15^ and that TNF induces endocytosis of occludin on epithelial cells^34^, we investigated whether occludin localization was altered in TNF^ΔARE/+^ mice. Surprisingly, we found no difference in occludin localization between TNF^ΔARE/+^ mice and WT littermates at 6 weeks of age (Supplementary Figure 3d), indicating that this decreased motility was not due to reduced interactions with epithelial occludin. We recently reported that reduced track speed and increased dwell time are key features associated with the ability of γδ IELs to facilitate the extrusion of apoptotic enterocytes in response to pathological concentrations of TNF^35^. Despite an increase in epithelial cell shedding in 6-week-old TNF^ΔARE/+^ mice (Supplementary Figure 3e), we were not able to visualize γδ IEL interactions with shedding epithelial cells *in vivo*, which may be due to the substantial reduction in γδ IEL number and motility. We conclude that fewer γδ IELs are relegated to cover a comparable area of epithelium at a slower speed, leading to compromised surveillance of the barrier prior to the onset of inflammation.

### TNF^ΔARE/+^ mice exhibit an early reduction in the frequency of immunosuppressive ileal CD39^+^ γδ IELs

During the pathogenesis of CD-like ileitis in TNF^ΔARE/+^ mice, we observed that mucosal homeostasis is disrupted; a loss of innate-like and regulatory IELs is followed by an expansion of induced IELs within the epithelium (Figure 1e-i). CD39, a cell-surface ectonucleosidase, works in concert with CD73 to convert adenosine triphosphate (ATP) to adenosine, an immunosuppressive molecule that dampens effector T cell responses^36^. While CD39 is often associated with conventional Foxp3^+^ Tregs, CD39^+^ γδ T cells exhibit immunosuppressive functions in the skin, lymph nodes and within tumors^37,38^. Recent reports indicate a reduction in the frequency of CD39^+^ γδ IELs in biopsies from pediatric IBD patients^39^, thus we hypothesized that CD39^+^ γδ IELs might be dysregulated in the events leading up to chronic ileitis.

Flow cytometric analysis revealed a substantial decrease in CD39^+^ γδ IELs across all Vγ subsets from as early as 4 weeks of age (Figure 4a,b); however, the relative expression of CD39 remained consistent (Figure S4a). Whereas nearly all γδ IELs express CD73 (Figure S4b), its expression is reduced in the majority of Vγ subsets at these early timepoints (Figure 4c,d). To determine whether CD39^+^ γδ IELs function in an immunoregulatory capacity, we performed suppression assays using sorted CD39^+^ or CD39^-^γδ IELs treated with extracellular (e)ATP in the presence or absence of a pharmacological CD39 inhibitor (POM-1)(Figure 4e). Culture with supernatant generated from CD39^+^ γδ IELs was sufficient to reduce the frequency of TNF^+^ CD8^+^ T cells, but only in the presence of eATP (Figure 4e,f). The addition of POM-1 abrogated this response, and culture with supernatant from CD39-deficient γδ IELs had no effect, indicating that CD39 expression and/or activity is required for immunosuppression. Interestingly, we did not observe a reduction in IFNγ production or cell proliferation (Figure S5a-c). Previous reports showed that regulatory CD39^+^ γδ T cell populations express conventional Treg markers such as Foxp3 and/or CD25^37,40^, yet expression of both markers was low in CD39^+^ γδ IELs (Figure S5d,e). In summary, we find that γδ IEL effector function is impaired before the observed reduction in γδ IEL number in TNF^ΔARE/+^ mice, suggesting that defective regulatory capacity of these cells may contribute to the initiation of ileitis.

**Figure 4:**
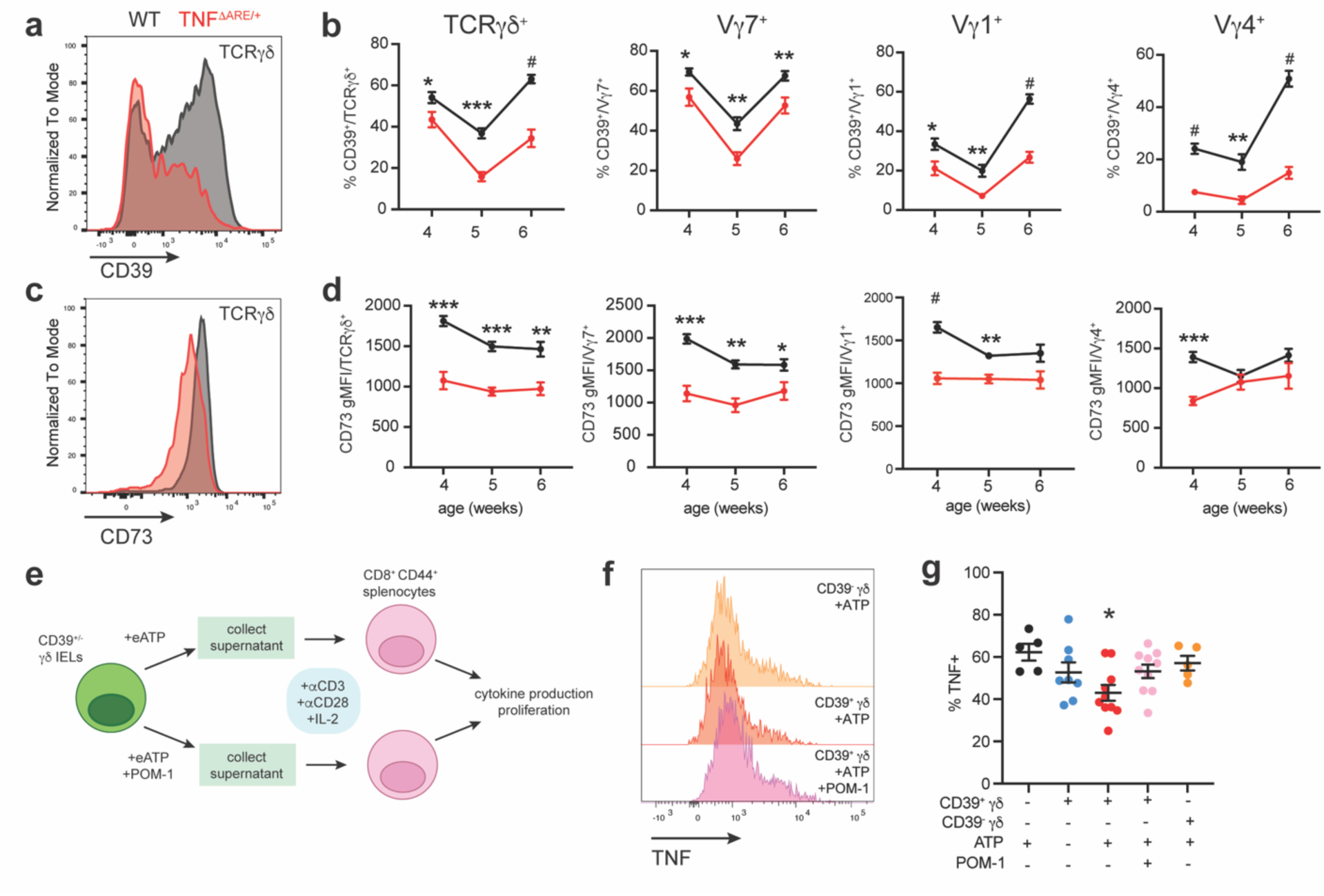
The frequency of immunoregulatory CD39* γδ IELs is decreased in TNF^ΔARE/+^ weanling mice. (a) Histogram of CD39 expression on ileal γδ IELs of WT and TNF^ΔARE/+^ mice at 6 weeks of age. (b) Frequency of CD39+ Vγ IEL subsets in 4-6-week-old WT and TNF^ΔARE/+^ mice, (c) Histogram of CD73 expression on ileal γδ IELs isolated from 4-week-old WT and TNF^ΔARE/+^ mice. (d) Geometric mean fluorescence intensity (gMFI) of CD73 in Vγ IEL subsets in 4-6-week-old WT and TNF^ΔARE/+^ mice, n = 4-11. Each data point represents an individual mouse, (e) Schematic outlining the experimental design of suppression assays, (f) Histogram of TNF expression in CD8+ CD44+ memory T cells following culture with supernatant from CD39+ or CD39-γδ IELs in the presence of eATP or POM-1. (g) Frequency of TNF+ CD8+ CD44+ memory T cells. n=5-10 experimental replicates. All data are shown as mean ± SEM from at least 2 independent experiments. Statistical analysis: b,d: unpaired t-test; g: one-way ANOVA, Tukey’s posthoc. *P<0.05, **P<0.01, ***p< 0.001, #P< 0.0001.

### Peripheral γδ T cells infiltrate the IEL compartment prior to disease onset

Our finding that γδ IEL regulatory function is compromised as early as 4 weeks of age led us to investigate whether additional cell surface markers associated with IEL functionality were similarly altered^28^. IELs are phenotypically characterized as CD44^+^ CD8α^+^ CD5^-^, which is reflective of their activated state and adaptation to the epithelial microenvironment^41–43^. Assessment of these cell surface receptors, in conjunction with CD39 and CD73, revealed that the frequency of those IELs that were CD8α^+^ and CD44^+^ was reduced among CD39^-^ γδ IELs relative to CD39^+^ γδ IELs (Figure 5a,b). Conversely, CD5 expression was increased. These observations were also reflected across individual Vγ IEL subsets (Figure S4b, S6). When compared to γδ IELs, fewer γδ T cells in mesenteric lymph nodes (MLN) expressed CD8α or CD44, and instead expressed CD5 (Figure 5c). Since the number of Vγ1^+^ and Vγ4^+^ IELs is increased in the absence of a proliferative response (Figure 2h,i, S2c,d), we hypothesized that these CD39^-^ γδ T cells were recent emigrants from the periphery. At 6 weeks of age, we detected an expansion of CD3^+^ T cells, including Vγ1^+^ and Vγ4^+^ T cells, in the MLN of TNF^ΔARE/+^ mice compared to WT (Figure S7, Figure 5d). Moreover, the frequency of CD103^+^ Vγ1^+^ and Vγ4^+^ T cells was increased in the MLN of weanling TNF^ΔARE/+^ mice (Figure 5e,f), indicating enhanced trafficking of peripheral γδ T cells to the gut. These data suggest that CD103^+^ CD39^-^ Vγ1^+^ and Vγ4^+^ T cells likely fail to upregulate CD39 and other markers associated with epithelial residency following migration into the gut. Thus, we conclude that reshaping of the γδ IEL compartment prior to the onset of ileal inflammation occurs when an influx of peripheral γδ T cells into the epithelium is unable to compensate for the significant loss of immunosuppressive tissue-resident γδ IELs.

**Figure 5:**
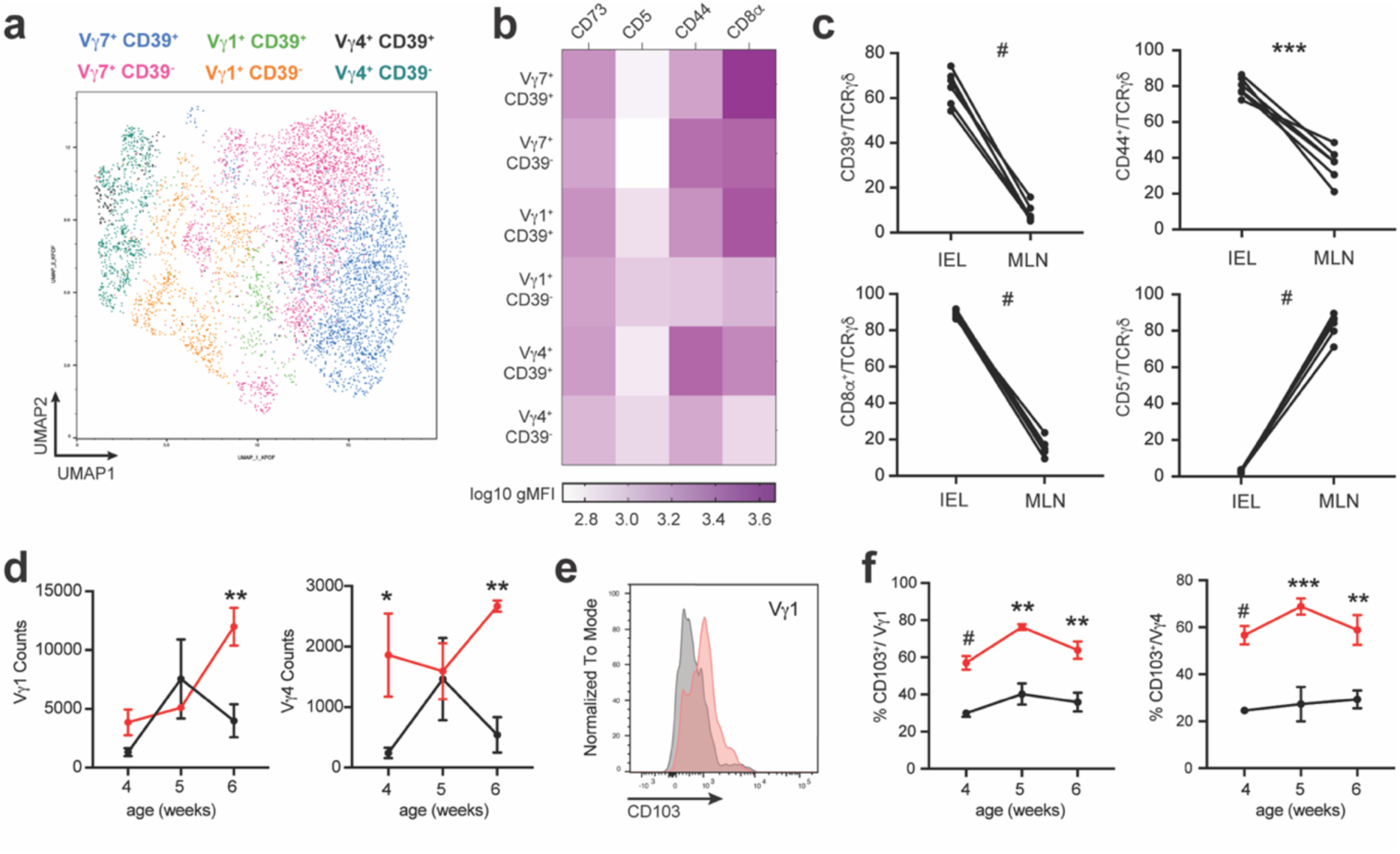
Peripheral Vγ1+ and Vγ4+ T cells migrate into the IEL compartment in TNF^ΔARE/+^ mice and fail to upregulate markers of tissue residency. (a) UMAP visualization of CD39-positive and -negative ileal Vγ IEL subsets and (b) heatmap showing CD73, CD5, CD44 and CD8α expression within these subsets in 6-week-old WT and TNF^ΔARE/+^ mice, (c) Comparison of the frequency of CD39+, CD44+, CD8α+ or CD5+ γδ T cells in the IEL compartment and MLN in 6-week-old WT mice, (d) Total number of Vγ1+ and Vγ4+ T cells in MLN of 4-6-week-old WT and TNF^ΔARE/+^ mice, (e) Representative histogram of CD103 expression in Vγ1+ T cells from the MLN of a 4-week-old WT or TNF^ΔARE/+^ mouse, (f) Frequency of CD103+ Vγ1+ and Vγ4+ T cells isolated from the MLN of WT orTNF^ΔARE/+^ mice, n = 4-11. All data are shown as mean ± SEM from at least 2 independent experiments. Each data point represents an individual mouse. Statistical analysis: c, paired t-test; d,f: unpaired t-test. *P<0.05, **P<0.01, *“P< 0.001, #P< 0.0001.

### Early depletion of the γδ IEL compartment correlates with increased severity of CD-like ileitis

Next, we inducibly depleted γδ T cells at 4 weeks of age using TcrdGDL; TNF^ΔARE/+^ mice to determine whether premature loss of all γδ T cells affected disease outcome. To our surprise, we observed a strain-dependent increase in the severity of ileal inflammation in 6-week-old, untreated TcrdGDL; TNF^ΔARE/+^ mice (Figure 6a,b), which was not further enhanced by diphtheria toxin (DTx) administration. To address the potential cause underlying the accelerated timeline of disease onset, we assessed γδ IEL number by *in situ* hybridization in both strains of TNF^ΔARE/+^ mice and found that the line harboring the TcrdGDL allele had reduced frequency of γδ IELs compared to the parental (P) strain, both P-TNF^ΔARE/+^ mice and P-WT controls (Figure 6c,d). We confirmed these findings with GFP immunostaining in mice on the TcrdGDL background, and again found that γδ IEL number at steady-state was reduced compared to 6-week-old P-TcrdGDL mice (Supplementary Figure 8a,b). Consistent with this, *Btnl1* expression was significantly reduced in 6-week-old TcrdGDL; TNF^ΔARE/+^ mice compared to P-TNF^ΔARE/+^, with a similar trend toward decreased *Hnf4g* expression (Figure 6e,f). Moreover, we detected an increase in TNF transcripts in TcrdGDL; TNF^ΔARE/+^ mice, whereas there was no difference in *Tnf* expression between 6- and 8-week-old P-TNF^ΔARE/+^ mice (Figure 6g). A recent report highlighted a correlation between segmented filamentous bacteria (SFB) and the severity of chronic ileitis in the TNF^ΔARE/+^ mouse model^44^; therefore, we next assessed relative SFB colonization in fecal samples from these mice. While we detected SFB in P-TcrdGDL mice, the abundance was higher in this line than in either P-TNF^ΔARE/+^ or TcrdGDL; TNF^ΔARE/+^ lines (Supplementary Figure 8c,d). Further, we did not observe differences in fecal SFB levels or IL-17 between the two lines (Supplementary Figure 8d, Figure 6h). In contrast to previous findings^44^, we failed to detect an increase in SFB during disease progression in TNF^ΔARE/+^ mice in our facility (Supplementary Figure 8e). Although the underlying cause of accelerated disease has yet to be determined, these data suggest that increased TNF expression and reduced γδ IEL number may contribute to a more rapid onset of ileal inflammation. Taken together, these studies support our findings that the early loss of γδ IELs correlates with increased susceptibility to CD-like ileitis; however, it remains unclear as to whether loss of γδ IELs is an initiating event or constitutes one of a series of events leading to disease pathogenesis.

**Figure 6.**
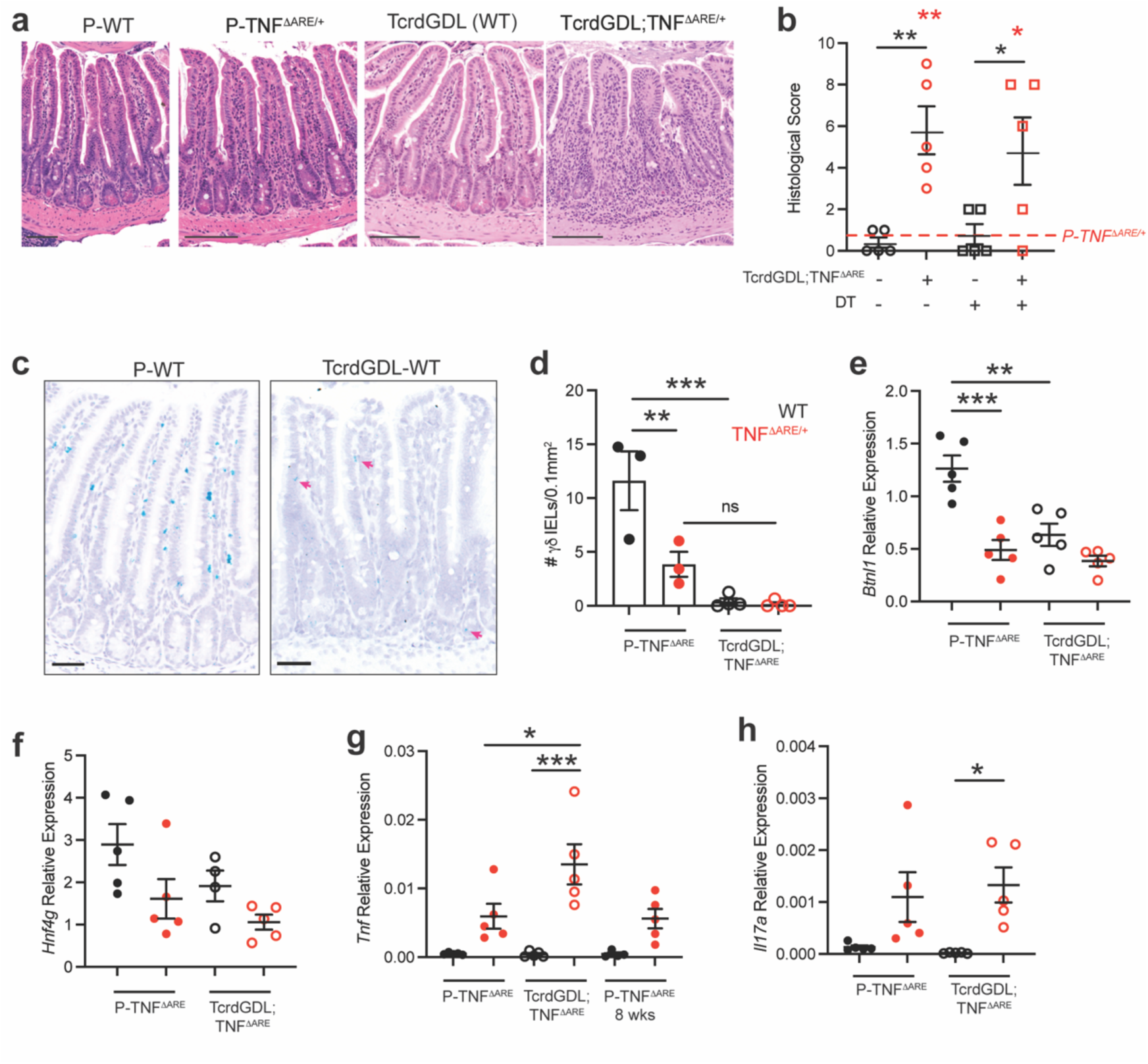
Earlier depletion of γδ IELs correlates with accelerated onset of ileal inflammation. (a) Representative H&E micrographs and (b) histological scores of ilea from 6-week-old WT and TNF^ΔARE/+^ mice of the parental (P) strain and those crossed to the TcrdGDL line. TcrdGDL; TNF^ΔARE/+^ mice received either PBS or diphtheria toxin (DT) at 4 weeks of age. Scale bar = 100 µm. Red asterisks indicate significance compared to P-TNF^ΔARE/+^ mice, (c) In situ hybridization for *Trdc* in ileal sections from 6-week-old P-WT or TcrdGDL-WT mice. Scale bar = 50 µm. (d) Morphometric analysis of γδ IELs in 6-week-old P-WT and p-TNE^ΔARE/+^ mice compared to those crossed to TcrdGDL line. Relative expression of (e) *Btnll,* (f) *Hnf4g,* (g) *Tnfand* (h) *II17a* in ileal tissues from mice described above. n=3-5. All data are shown as mean ± SEM from at least 2 independent experiments. Each data point represents an individual mouse. Statistical analysis: one-way ANOVA with Sidak’s posthoc test *P<0.05, **P<0.01, ***P< 0.001.

## Discussion

In this study, we provide the first evidence that γδ IELs are dysregulated prior to the onset of chronic ileitis. Analysis of the IEL compartment of TNF^ΔARE/+^ mice revealed that immunosuppressive CD39^+^ γδ IELs are reduced starting at 4 weeks of age, with a decrease in total γδ IEL number and frequency apparent one week later. We identify a simultaneous increase in CD103^+^ CD39^-^ peripheral γδ T cells within the epithelium, yet these cells are unable to functionally compensate for the loss of the CD39^+^ γδ IEL population. Thus, we posit that impaired γδ IEL regulatory function may result in the loss of mucosal tolerance as indicated by the expansion of induced IELs during disease progression. Further, we observed that mice with a diminished γδ IEL compartment exhibit accelerated disease development, demonstrating a protective role of γδ IELs in the context of ileal inflammation.

Although decreased γδ IEL frequency has been reported in ileal CD^16^, the molecular mechanisms involved in the disruption of γδ IEL homeostasis during chronic intestinal inflammation remain unclear. Samples obtained from patients with active IBD exhibit reduced *BTNL8* expression, the human homolog to murine *Btnl6*^45^. Moreover, a polymorphism in *Hnf4a*, a transcription factor which regulates the expression of epithelial BTNL proteins^29^, is associated with increased IBD susceptibility^46^. We find that *Hnf4g* expression is reduced in the ileal epithelium of TNF^ΔARE/+^ mice at 4 weeks of age, leading to a subsequent reduction in *Btnl1/6* that coincides with a loss of γδ IEL number and a shift in the γδ IEL compartment toward Vγ7^-^ subsets. Although HNF4A and HNF4G are thought to exhibit a degree of functional redundancy^29^, we did not observe a change in *Hnf4a* expression, which may be attributed to regional specification of these proteins. These data indicate that defects in the HNF4G/BTNL axis adversely affect the γδ IEL compartment prior to the onset of chronic ileitis. It remains possible that alteration of other epithelial- or microbial-derived signals contribute to γδ IEL depletion during disease initiation. Although IL-15 is required for γδ IEL homeostasis^47–49^, increased IL-15 expression was reported in TNF^ΔARE/+^ mice^50^ and IBD patients^51^ suggesting that the loss of γδ IELs occurs independent of IL-15. While aryl hydrocarbon receptor (AhR) signaling is implicated in both γδ IEL maintenance and intestinal inflammation^52–54^, further study is needed to determine whether defects in this pathway contributes to γδ IEL depletion at these early timepoints.

Of the remaining γδ IELs, we observed a stark reduction in γδ IEL surveillance at 6 weeks of age in TNF^ΔARE/+^ mice. We initially hypothesized that TNF-induced internalization of occludin would limit γδ IEL motility^15^ yet we were unable to discern reorganization of epithelial occludin early in disease development, consistent with previous reports^55^. Development of co-culture models to assess γδ IEL interactions with CD patient-derived enteroids may shed light on the underlying mechanisms by which γδ IEL motility is inhibited. In addition to immunosurveillance, we recently showed that γδ IELs facilitate the shedding of apoptotic enterocytes in response to TNF exposure^35^. Although we found increased epithelial apoptosis in 6-week-old TNF^ΔARE/+^ mice, the reduced number of γδ IELs limited our ability to observe shedding-associated migratory behaviors. This impaired surveillance has negative implications for limiting microbial translocation^14^, and may also compromise potential γδ IEL interactions with specialized epithelial cells along the crypt-villus axis. For example, γδ IELs were recently shown to promote Paneth cell survival through secretion of the anti-apoptotic factor API5^27^. The loss of γδ IELs appears to precede Paneth cell loss in TNF^ΔARE/+^ mice, which typically coincides with disease onset^56^. Thus, future studies will focus on the involvement of γδ IELs in Paneth cell homeostasis during the initiation and progression of chronic ileitis.

The role of γδ T cells in ileitis development has been ambiguous due to caveats regarding the experimental design of previous studies^17,18^. Antibody-mediated blockade of TCRγδ signaling starting at 4 weeks in TNF^ΔARE/+^ mice resulted in exacerbated ileitis at 14 weeks, indicating that inhibition of adaptive γδ IEL function increases susceptibility to ileal inflammation^57^. Despite our intentions to inducibly deplete γδ T cells in TNF^ΔARE/+^ mice to elucidate the requirement of both the innate and adaptive contributions of γδ IELs in chronic ileitis, we unexpectedly observed a reduction in γδ T cells in TcrdGDL; TNF^ΔARE/+^ mice at steady-state. Comparisons between this new line and the P-TNF^ΔARE/+^ line revealed enhanced TNF production early in life that did not correlate with SFB colonization. Based on the presence of an intact γδ IEL compartment in the P-TcrdGDL line, we posit that crossing these dams to P-TNF^ΔARE/+^ sires may have selected for a microbiota that accelerates disease development. Since the presence of gut bacteria is required for ileitis development^58^, it is challenging to directly test this hypothesis. Despite not knowing the precise cause of accelerated ileitis in this model, the correlation between a more rapid depletion of γδ IELs and disease development, combined with reduced γδ IEL number prior to the onset of inflammation in two models of ileitis, clearly demonstrates the protective role of γδ IELs in the maintenance of mucosal homeostasis.

Previous studies highlight an immunosuppressive role for colonic γδ IELs in experimental colitis largely thought to be dependent on TGFβ and IL-10 production^19,57,59^. In the small intestine, γδ IELs secrete minimal amounts of cytokine under homeostatic conditions^27,60^ suggesting that hydrolysis of ATP into adenosine is the most likely mechanism by which these cells exert their regulatory function. While murine peripheral CD39^+^ γδ T cells suppressed naïve CD4^+^ T cells via IL-10^37^, another group found that adenosine, and not IL-10 or TGFβ, contributed to the regulatory activity of human tumor infiltrating CD39^+^ γδ Tregs^38^. We now demonstrate that CD39^+^ γδ IELs inhibit effector cytokine production through an adenosine-dependent mechanism; however, the specific contribution of CD39^+^ γδ IELs to disease pathogenesis and their influence on local effector lymphocytes remains an exciting area for future investigation.

Genetic polymorphisms identified in the promoter region of CD39, which result in reduced gene expression, are associated with IBD^61^. Single cell RNAseq analysis of biopsies from CD patients revealed both a decrease in γδ IELs and a reduction in those expressing CD39 compared to healthy controls^16^. The frequency of CD39^+^ γδ IELs is similarly reduced in patients with colonic CD yet this does not correlate with disease severity^62,63^. In support of its regulatory function, increased CD39 expression was observed in IBD patients who responded to anti-TNF therapy^64^. A recent study demonstrated that treatment with dipyridamole, which prevents depletion of intracellular cAMP and can increase CD39 expression on lymphocytes, ameliorated experimental colitis and improved clinical, endoscopic and histological parameters in pediatric colitis patients^39^. Taken together, these findings suggest that the enhancement of adenosine-mediated suppressive capacity of γδ IELs may provide a viable therapeutic strategy to promote mucosal tolerance and dampen pro-inflammatory responses within the IEL compartment.

We were surprised to find an increase in Vγ1^+^ and Vγ4^+^ IELs in TNF^ΔARE/+^ mice at 6 weeks of age, especially in the absence of local proliferation within the epithelium. However, the cell surface receptor profile of these IELs indicates that these Vγ1^+^ and Vγ4^+^ cells recently emigrated from the periphery to the intestinal epithelium. Failure to upregulate receptors commonly associated with natural IELs suggests that these Vγ7^-^ cells may lack the appropriate environmental cues to promote maturation within the epithelial compartment, and thus limit their ability to adopt a CD39^+^ regulatory phenotype. Additional studies are needed to determine (1) the extent to which the epithelial signalome is altered at these early timepoints, (2) whether these peripheral emigrants actively contribute to disease pathogenesis, and (3) if the loss of CD39^+^ γδ IELs is reversible following disease onset. Previous work demonstrated that the loss of BTNL expression and an influx of peripheral IFNγ-producing Vδ1^+^ cells permanently reshaped the γδ IEL compartment prior to inflammation in celiac disease patients^65^, yet whether this also occurs in the context of IBD remains unknown.

Based on our findings, we assert that restoring both the number and function of γδ IELs may be key to preventing the onset or relapse of ileitis. Our laboratory has previously discovered a transmissible microbiota that enhances the migratory and proliferative capacity of γδ IELs^66^. Further studies will address whether transfer of this hyperproliferative γδ IEL-associated microbiota into TNF^ΔARE/+^ mice is sufficient to augment or restore γδ IEL number and functionality, and to mitigate the development or severity of ileal pathology. Moreover, assessment of γδ IEL number or CD39 expression on these cells may serve as a useful biomarker to identify potential relapse or predict responsiveness to therapy.

## Materials and Methods

### Animals

Mice of both sexes were used between 4-17 weeks of age and maintained on a C57BL/6 background under specific pathogen-free (SPF) conditions in Allentown caging with aspen shavings and LabDiet5010 chow (Purina, St. Louis, MO) and RO water. TNF^ΔARE/+^ mice^8^ were rederived by the Genome Editing Shared Resource, Rutgers Cancer Institute of New Jersey and subsequently crossed to TcrdGDL mice^67^. TNF^+/+^ littermates were used as controls. For suppression assays, wildtype C57BL/6 mice or TcrdEGFP^68^ mice were used between 8-14 weeks of age. To assess IEL proliferation, 200 μg of 5-ethynyl-2′-deoxyuridine (EdU)(Sigma-Aldrich) was administered i.p. daily for 4 days. To induce γδ T cell depletion in 4-week-old TcrdGDL; TNF^ΔARE/+^ mice, 15 ng of diphtheria toxin (List Biological, Campbell, CA) or endotoxin-free PBS per gram body weight was administered i.p. 24 and 48 h prior to the experiment. Mouse weights and clinical scores (0 – normal, 1 – abnormal, 2 – severe for each of the following categories: fur texture, posture, activity) were recorded. All studies were conducted in an Association of the Assessment and Accreditation of Laboratory Animal Care-accredited facility according to protocols approved by Rutgers New Jersey Medical School Comparative Medicine Resources, Case Western Reserve University and the Icahn School of Medicine Center for Comparative Medicine and Surgery.

### Histology

Swiss-rolled small intestine was fixed in 10% neutral buffered formalin, paraffin-embedded, sectioned at 5 μm and stained with hematoxylin and eosin. Total histological score was calculated, blinded to genotype and condition, based on the following criteria: immune cell infiltration (0 – no increase in LP; 1 – mild increase in LP; 2 – moderate increase in LP; 3 – strong increase in LP; 4 – mild increase in submucosa; 5 – moderate increase in submucosa; 6 – strong increase in submucosa), villous damage (0 – normal; 1 – blunted (drumstick appearance); 2 – shortening of villi; 3 – complete destruction of villi), submucosal inflammation (0 – 0% involvement; 1 – 1-25% involvement; 2 – 25-50% involvement; 3 – 50-75% involvement; 4 – 75-100% involvement), and transmural inflammation (0 – 0% involvement; 1 – 1-25% involvement; 2 – 25-50% involvement; 3 – 50-75%; 4 – 75-100%).

### Flow Cytometry

Ileal IELs were isolated as previously described^15^ and mesenteric lymph nodes (MLN) from each mouse were collected. Spleens were homogenized using a 100 μm filter (CellTreat) and red blood cells were lysed. Lymphocytes were stained using fixable viability dye eFluor 780 (eBioscience) or Zombie UV (Biolegend), anti-CD3 (145-2C11, BD Biosciences and BioLegend), anti-TCRγδ (GL3, BioLegend), anti-TCRβ (H57-597, BioLegend), anti-CD4 (RM4-5, Biolegend), anti-CD8α (53-6.7, Biolegend), anti-CD8β (YTS1567.7, Biolegend), anti-Vγ1 (2.11, BD Biosciences), anti-Vγ4 (UC3-10A6, BD Biosciences), anti-Vγ7 (GL7, provided by Rebecca O’Brien (National Jewish Health, Denver, CO)), streptavidin-BUV737 (BD Biosciences), anti-Vδ4 (GL2, BD Biosciences), anti-CD39 (Duha59, Biolegend), anti-CD73 (TY/11.8, Biolegend), anti-CD44 (1M7, Biolegend), anti-CD5 (53.7-3, Biolegend), anti-CD103 (2E7, Biolegend), anti-CD25 (PC61, Biolegend), Apotracker green (Biolegend), Click-iT Plus EdU Alexa Fluor 647 Flow Cytometry Assay Kit (Invitrogen) and Brilliant Stain Buffer (BD Biosciences). Precision Count Beads (Biolegend) were used to generate absolute counts of MLN cells. For intracellular staining, samples were either fixed with Cytofix/Cytoperm (Becton Dickinson) prior to staining with anti-IFNγ (XMG1.2, Biolegend) or anti-TNFα (MP6-XT22, BD) or eBioscience Foxp3 transcription factor kit followed by staining with anti-Foxp3 (FJK-16s, ThermoScientific). Flow cytometry was performed on a LSR Fortessa (BD Biosciences, Franklin Lakes, NJ) and the data analyzed using FlowJo (v. 10.8.2; Tree Star).

### Immunostaining

For immunofluorescence staining, 7 μm thick frozen sections of ileal tissue were blocked in 10% NGS and stained with biotinylated anti-GFP antibody (Abcam, ab6658, 1:250) or anti-occludin AlexaFluor 594 (Invitrogen, OC-3F10, 1:100). Slides were incubated with streptavidin-AF488 (Jackson ImmunoResearch, 016-540-084, 1:500) or phalloidin AlexaFluor 647 (Life Technologies, 1:500) and Hoechst 33342 (Life Technologies, 1:1000) before mounting with Prolong Glass (Life Technologies). Fluorescent images were captured on an inverted DMi8 microscope (Leica Microsystems) equipped with a CSU-W1 spinning disk, ZYLA SL150 sCMOS camera (Andor), PL APO ×40/0.85 dry objective (Andor). The number of γδ IELs in ileal sections was quantified using FIJI (NIH).

For immunohistochemistry, antigen retrieval was performed using citric acid–based antigen unmasking solution (Vector Laboratories) on 5 μm sections from FFPE Swiss rolls of distal small intestine. Sections were subsequently blocked in 10% normal goat serum, incubated with rabbit anti-CC3 (5A1E, Cell Signaling) followed by horseradish peroxidase–conjugated anti-rabbit IgG (Vector Laboratories), developed using a 3,3′-diaminobenzidine solution, counterstained with hematoxylin, and mounted in Cytoseal XYL (ThermoScientific). Brightfield images were acquired using a BZ-X710 with x4/0.1 PL APO objective (Keyence). The number of CC3^+^ cells were quantified per mm of basement membrane.

### Quantitative real-time PCR

One cm pieces of terminal ileum were homogenized in TRizol (Life Technologies) using a bead beater (Mini-Beadbeater-24, BioSpec Products). Fresh fecal pellets were collected and stored at -80°C before processing. DNA was extracted using QIamp Fast DNA Stool Mini Kit following manufacturer’s instructions (Qiagen). RNA was extracted and purified using a RNeasy mini kit (Qiagen), after which cDNA was generated using an iScript cDNA Synthesis kit (Bio-Rad). qPCR was performed using 50ng of cDNA and PowerUp SYBR Green Master Mix (Applied Biosystems) on a QuantStudio *3/*6 Real-Time PCR System. Murine gene expression was normalized to m*Cdh1* expression or m*GAPDH*. Bacterial gene expression was normalized to 18S rRNA. The following primers were used:

*mBtnl1* (forward, 5’TGACCAGGAGAAATCGAAGG 3’;

*mBtnl1* (reverse 5’CACCGAGCAGGACCAATAGT 3’);

*mBtnl6* (forward, 5’ GCACCTCTCTGGTGAAGGAG 3’);

*mBtnl6* (reverse, 5’ ACCGTCTTCTGGACCTTTGA 3’);

*mHnf4a* (forward, 5’GTC CAG AGC TAG CGG AGA TG 3’);

*mHnf4a* (reverse, 5’ TAC TGC CGG TCG TTG ATG TA 3’);

*mHnf4g*(forward, 5’ GTG TCA ACT GTT TAT GTG CCA TC 3’);

*mHnf4g* (reverse, 5’GTT CAT TTT GCA CCG CTT CTT TT 3’);

*mCdh1* (forward, 5’ TCCTTGTTCGGATATGTGTC 3’);

*mCdh1* (reverse, 5’GGCATGCACCTAAGAATCAG 3’);

*mTnfa* (forward, 5’ CATCTTCTCAAAATTCGAGTGACAA 3’);

*mTnfa* (reverse, 5’ TGGGAGTAGACAAGGTACAACCC 3’);

*mIl17a* (forward, 5’ ACTACCTCAACCGTTCCACG 3’);

*mIl17a* (reverse, 5’ CTTCCCAGATCACAGAGGGA 3’);

*mGapdh* (forward, 5’ AAGGTGGTGAAGCAGGCATCTGAG 3’);

*mGapdh* (reverse, 5’ GGAAGAGTGGGAGTTGCTGTTGAAGTC 3’);

18S rRNA (forward, 5’ CATTCGAACGTCTGCCCTATC 3’);

18S rRNA (reverse, 5’ CCTGCTGCCTTCCTTGGA 3’);

SFB 16S rRNA (forward, 5’ GACGCTGAGGCATGAGAGCAT 3’);

SFB 16S rRNA (reverse, 5’ GACGGCACGGATTGTTATTCA 3’)

### In situ hybridization

RNA in situ hybridization was performed on 5 μm paraffin-embedded ileal sections using the RNAScope® 2.5 HD Duplex Manual Assay (Cat. no 322430) (Advanced Cell Diagnostics, Inc.) with the HybEZ™ Hybridization System per manufacturers’ instructions. RNAscope® probes used were: Mm-Btnl1 (Cat No. 436641), Mm-Btnl6-C2 (Cat No. 439821-C2), Mm-Trdc (Cat No. 449351) and 2-plex Negative Control Probe (Cat No. 320751). Spots were quantified using QuPath (v0.4.3).

### Intravital Microscopy

Intravital imaging was performed as previously described^69,70^. AlexaFluor 568 or AlexaFluor 647 hydrazide (Life Technologies) was used to visualize the intestinal lumen and Hoechst 33342 was used to label nuclei. 3D reconstructions of time-lapse videos were rendered by Imaris (v. 9.9, Bitplane) and an autoregressive tracking algorithm was employed to quantify IEL migration. Arrest coefficients were calculated based on the proportion of time in which an individual cell had an instantaneous speed of < 2 μm/min over the total length of the track^71^. Rose diagrams represent the total displacement of individual γδ IEL tracks.

### Suppression Assay

1.5×10^5^ CD39^+/-^ γδ IELs were sorted and cultured with IL15/15R complex (ThermoScientific, GRW15PLZ, 1 μg/ml) for one day. CD39^+^ γδ IELs were treated with 100 μM POM-1 (Biotechne) for 2 h prior to the addition of 10 μM ATP (ThermoScientific) and IL-2 (6 ng/ml, Peprotech). Supernatants were collected and stored at ^-^20°C for future use. 1×10^5^ splenic CD8^+^ CD44^+^ T cells were sorted, labeled with Cell Trace Violet (Invitrogen), stimulated with 1 μg/ml of plate-bound anti-CD3 (145-2C11, Biolegend), 2 μg/ml anti-CD28 (37.5, Biolegend) and IEL supernatant described above. On days 2 and 3, the culture supernatant was refreshed under the same treatment conditions. On day 4, following a 4 h treatment with Golgistop and Golgiplug (BD), cells were stained for analysis on day 4.

### Data Analysis

Statistical analysis was performed using Prism (GraphPad, version 9.5.1). Two-tailed unpaired student *t* tests were used to compare two independent samples, whereas a paired *t* test was used to compare samples from the same source. P≤ 0.05 was considered statistically significant. Comparisons between multiple independent variables were performed using one-way ANOVA followed by post-hoc Tukey or Šidák multiple comparisons tests. All data are presented as either the mean +/- SEM or with a 95% confidence interval.

## Acknowledgments

Cell sorting was performed at the Rutgers NJMS Flow Cytometry and Immunology Core Laboratory and supported by National Institute for Research Resources Grant (S10 RR027022). Ileal tissue samples from SAMP/AKR mice were provided by the Morphology/Imaging Core, Cleveland DDRCC (P30 DK097948) with sectioning performed by the Mount Sinai Biorepository and Pathology CoRE. This work was supported by A*STAR postdoctoral fellowship (W.X.), New Jersey Commission on Cancer Research COCR23PRF017 (N.B.G.), National Institute of Health Grants R21 AI143892, R21AI171959, and R01 DK135272, and Crohn’s and Colitis Foundation Senior Research Award (K.L.E.).

## Author Contributions

W.X. designed and performed experiments and wrote the manuscript. N.B.G. designed and performed experiments and wrote the manuscript. W.X. and N.B.G. contributed equally to the work. A.F. and S.A. performed experiments and analyzed the data. E.B.B. performed experiments. L.Z. analyzed data. M.R.F. provided resources and supervised the research. K.J.P., I.S., I.P., T.T.P., and G.K. provided resources. K.L.E. conceived the study, analyzed data, supervised the research and wrote the manuscript. All authors approved the final manuscript.

---

**Supplementary Figure 1:**
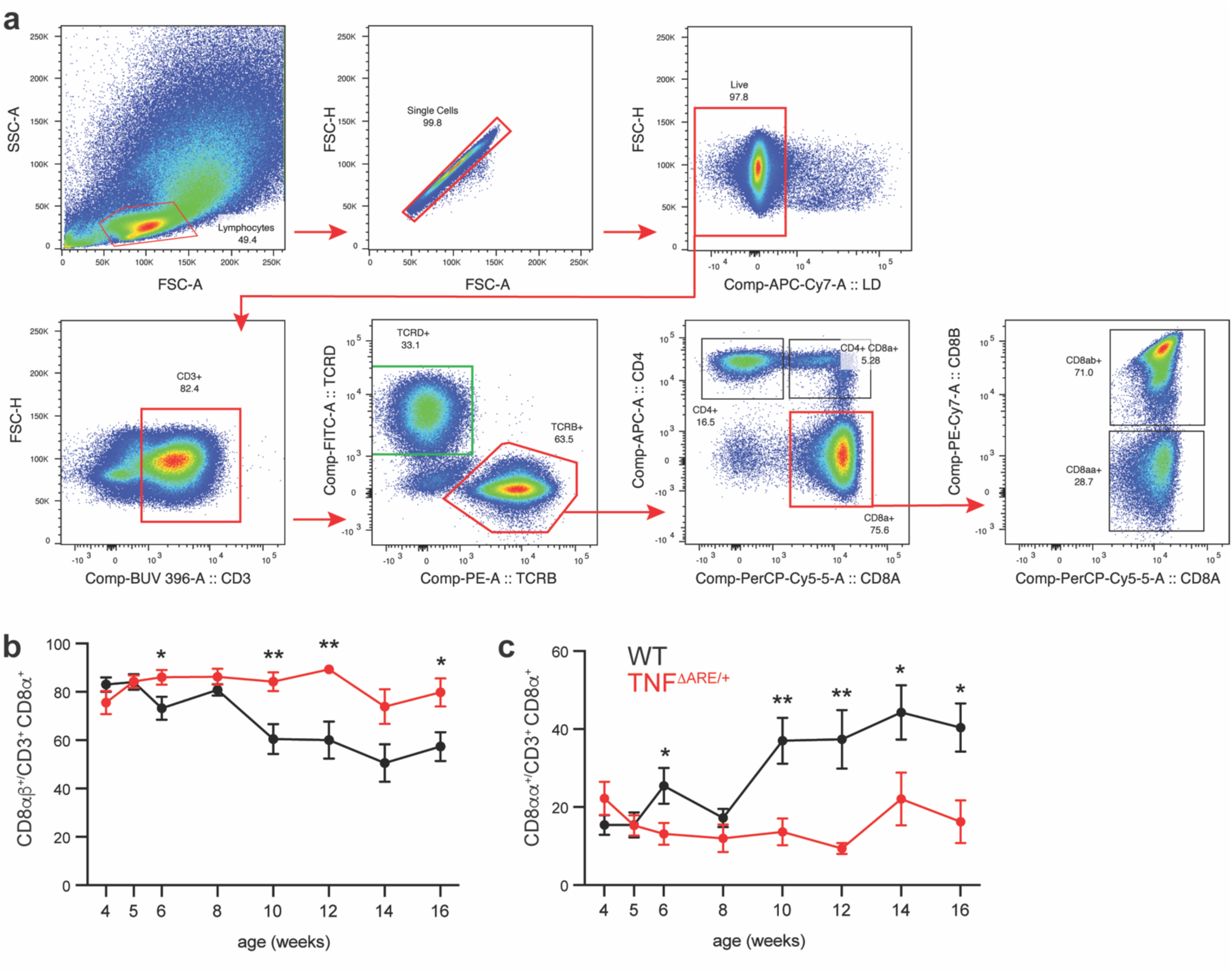
CD8α+ IELs are dysregulated following ileitis onset in TNF^ΔARE/+^ mice. (a) Gating strategy for ileal IEL subsets. Frequency of (b) CD8αα+ or CD8αβ IELs gated on CD3+ CD8α+ in WT or TNF^ΔARE/+^ mice from 4 to 16 weeks of age. n = 4.9. All data are shown as mean ± SEM from at least 2 independent experiments. Statistical analysis: unpaired t-test. *P<0.05, **P<0.01.

**Supplementary Figure 2:**
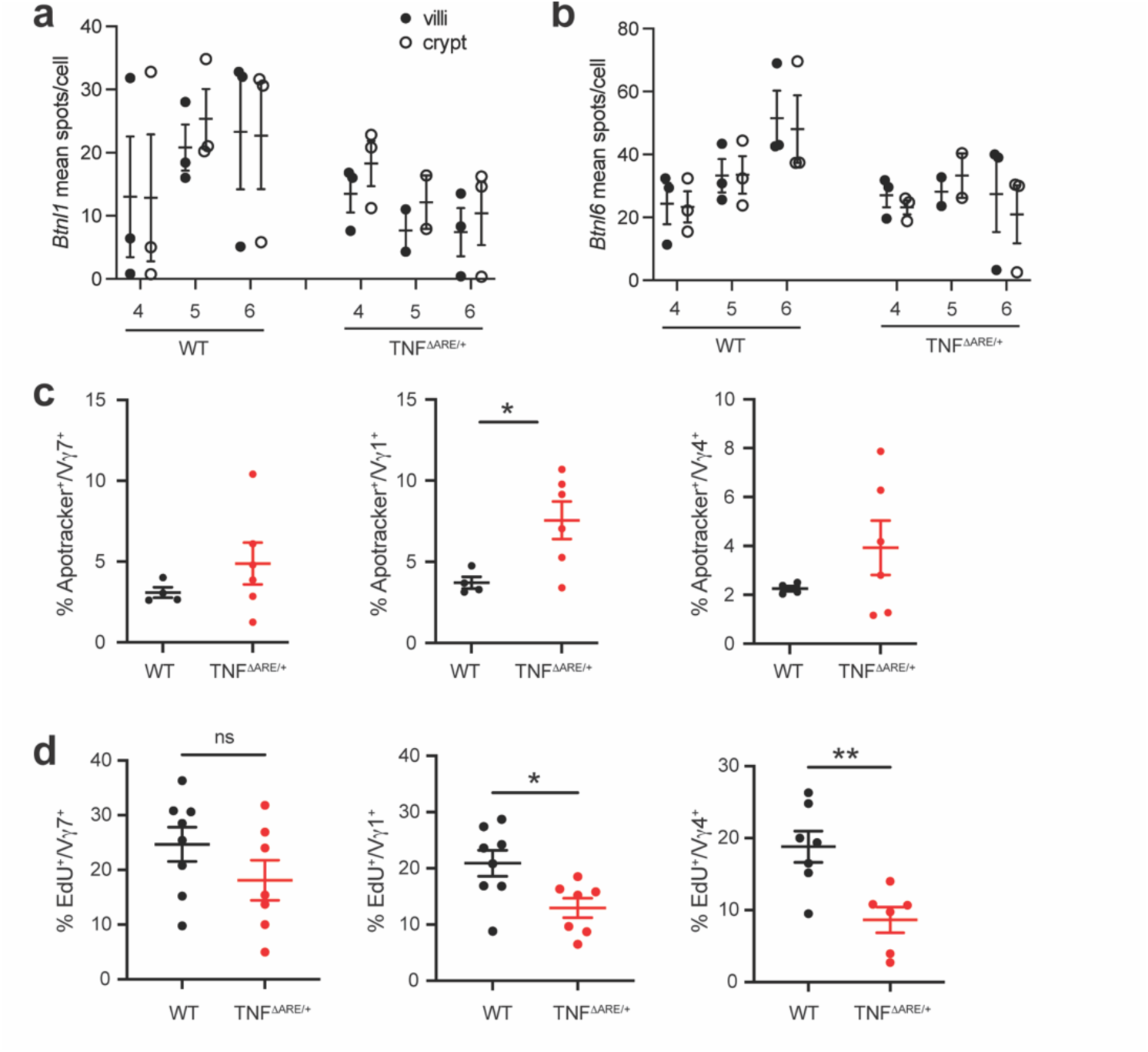
Increased apoptosis and proliferation defects are enhanced in Vγ7^-^ IEL subsets in young TNF^ΔARE/+^ mice. Quantification of (a) *Btnll* or (b) *Btnl6* spots/cell within ileal crypts or villi. n=2-3. Frequency of (c) EdU+ or (d) apoptotic Vγ7+, Vγ1+, Vγ4+ IEL subsets isolated from WT and TNF^ΔARE/+^ ilea at 5 weeks of age. n = 4-8. All data are shown as mean + SEM from at least 2 independent experiments. Each data point represents an individual mouse. Statistical analysis: unpaired t-test. *P<0.05, **P<0.01; ns not significant.

**Supplementary Figure 3.**
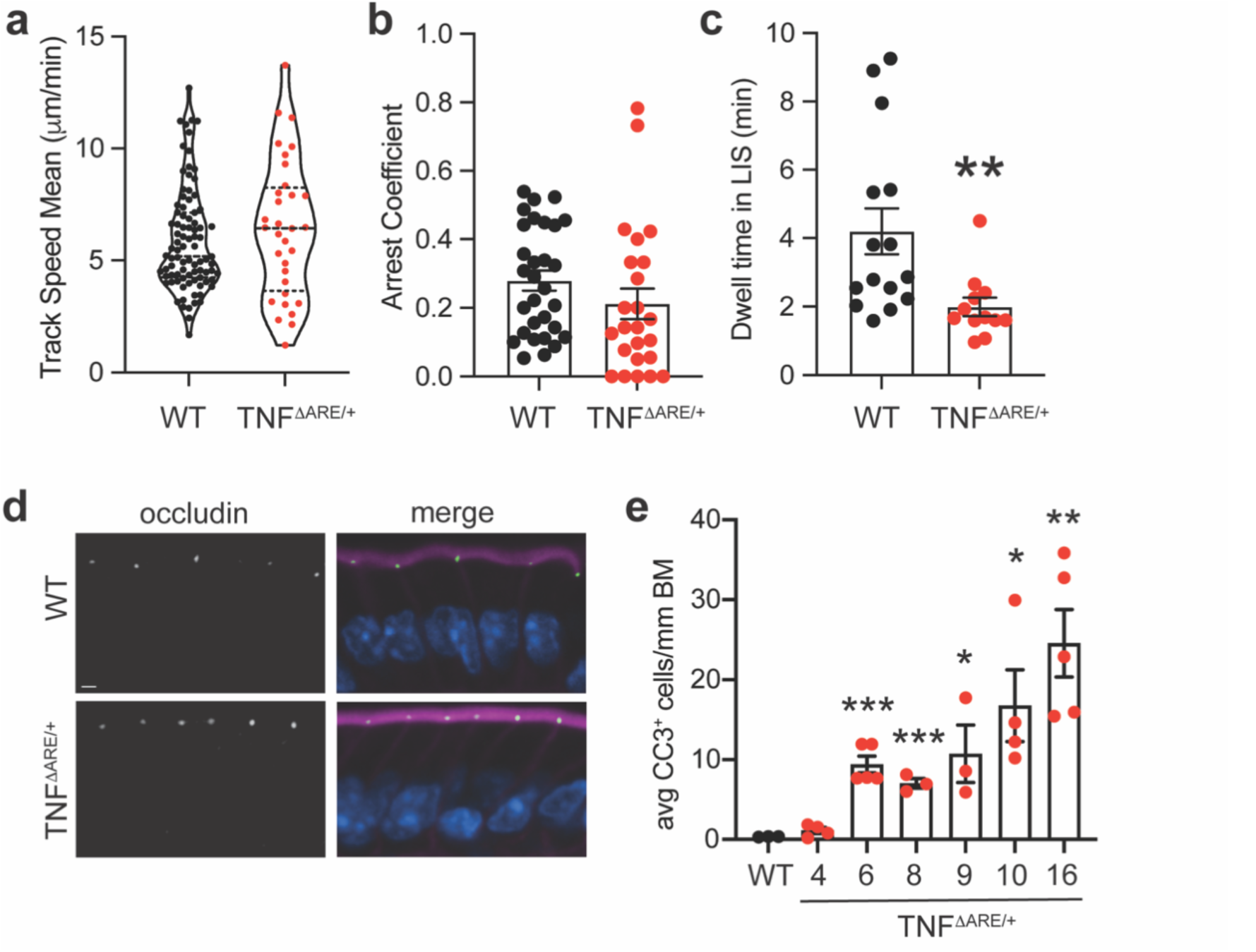
Impaired γδ IEL migratory behavior in TNF^ΔARE/+^ mice temporally coincides with increased epithelial cell apoptosis. Intravital microscopy was performed on ileal mucosa of 5-wk-old TcrdGDL; TNF+/+ (WT) or TcrdGDL; TNF^ΔARE/+^ mice, (a) Mean track speed, (b) arrest coefficient, and (c) dwell time of γδ lELs within the LIS. n= 88 tracks (WT), 32 tracks (TNF^ΔARE/+^). (d) Representative immunofluorescence micrograph of occludin (green) in ilea from 6-wk-old TcrdGDL TNF+/+ (WT) or TcrdGDL TNF^ΔARE/+^ mice. F-actin is shown in magenta and nuclei are blue. Scale bar = 3 µm. (e) Quantification of CC3+ enterocytes from WT (9 weeks of age) or TNF^ΔARE/+^ mice (4-16 weeks of age). n=3-5 mice. All data are shown as mean + SEM from at least 2 independent experiments. Each data point represents an individual track (a-c) or individual mouse (e). Statistical analysis: unpaired t-test; (e) compared to WT. *P<0.05, **P<0.01, ***P< 0.001.

**Supplementary Figure 4:**
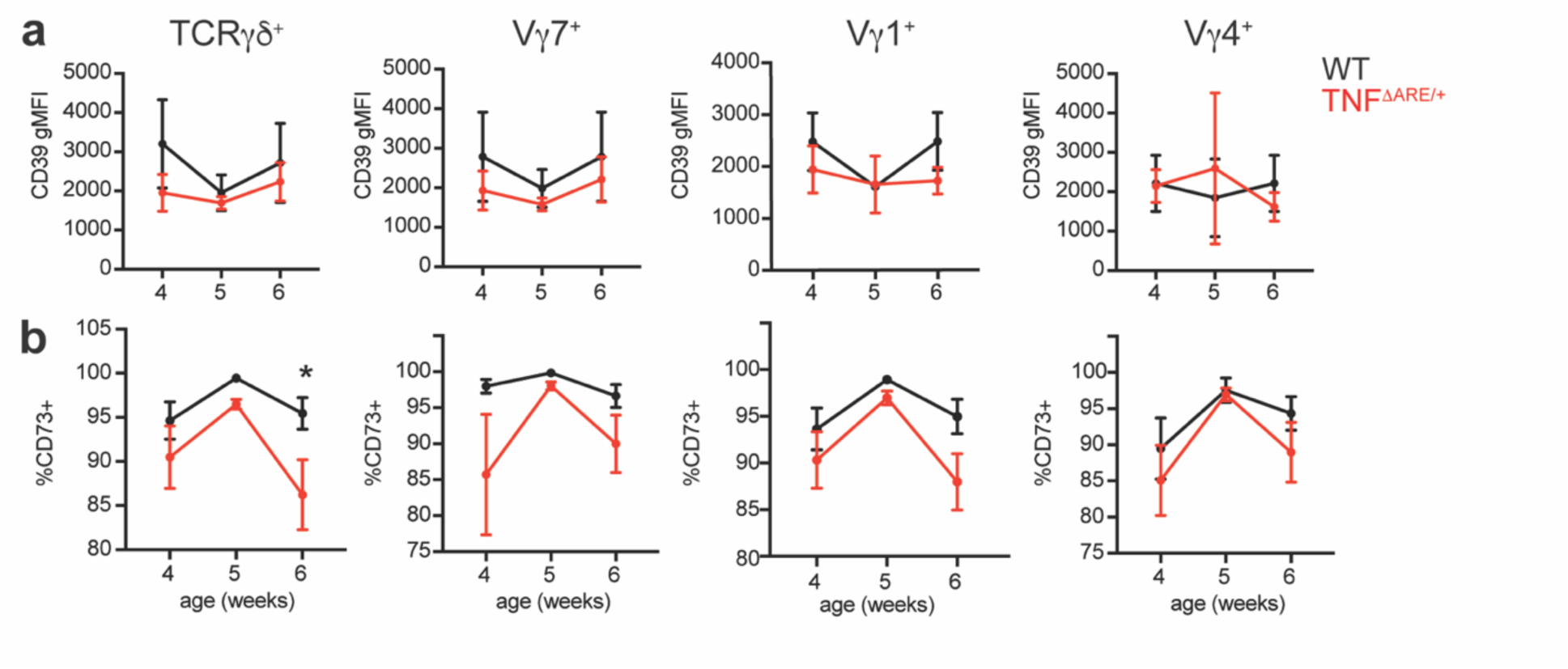
CD39 expression and the frequency of CD73-positivity is similar between Vγ IEL subsets in TNF^ΔARE/+^ weanling mice. (a) Geometric mean fluorescence intensity (gMFI) of CD39 expression in and (b) frequency of CD73+ Vγ IEL subsets in 4-6 week-old WT or TNF^ΔARE/+^ mice. n = 4-11. All data are shown as mean ± SEM from at least 2 independent experiments. Statistical analysis: unpaired t-test. *P<0.05.

**Supplementary Figure 5:**
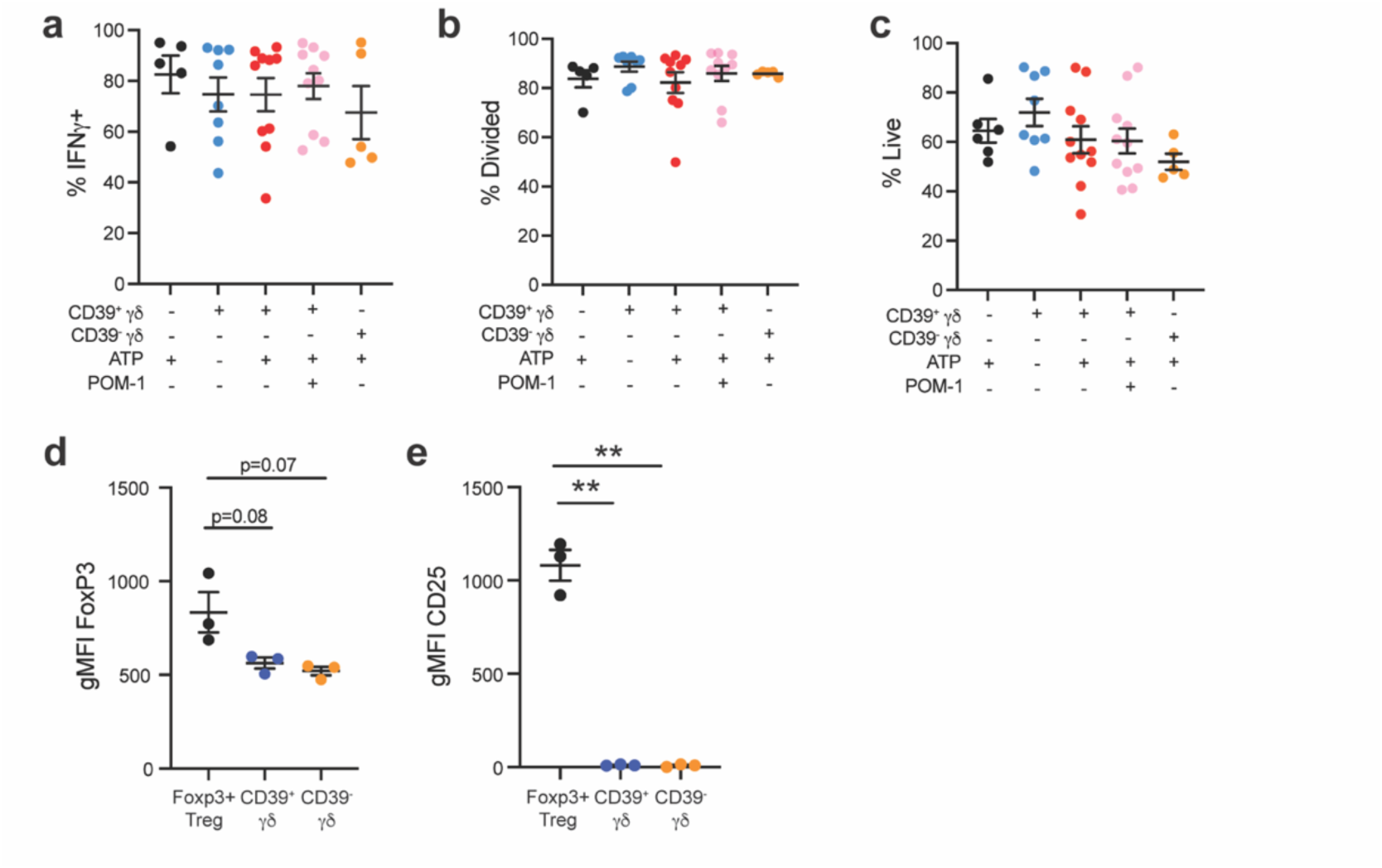
CD39* γδ T cells do not express conventional Treg markers or suppress CD8* proliferation or IFNγ production. CD8+ CD44+ memory T cells were stimulated and cultured with supernatant from CD39+ or CD39-γδ lELs in the presence of eATP or POM-1. The percentage of (a) IFNγ+ CD8+ T cells, (b) target cells that had divided, and (c) live target cells following culture are shown, n = 5-10 experimental replicates. The geometric mean fluorescence intensity (gMFI) of (d) FoxP3 and (e) CD25 expression in CD4+ Foxp3+ Tregs, CD39+ and CD39-γδ lELs. All data are shown as mean ± SEM from at least 2 independent experiments. n=3. Each data point represents an individual mouse. Statistical analysis: a-c: one-way ANOVA, Tukey’s posthoc; d,e: unpaired t-test. **P<0.01.

**Supplementary Figure 6:**
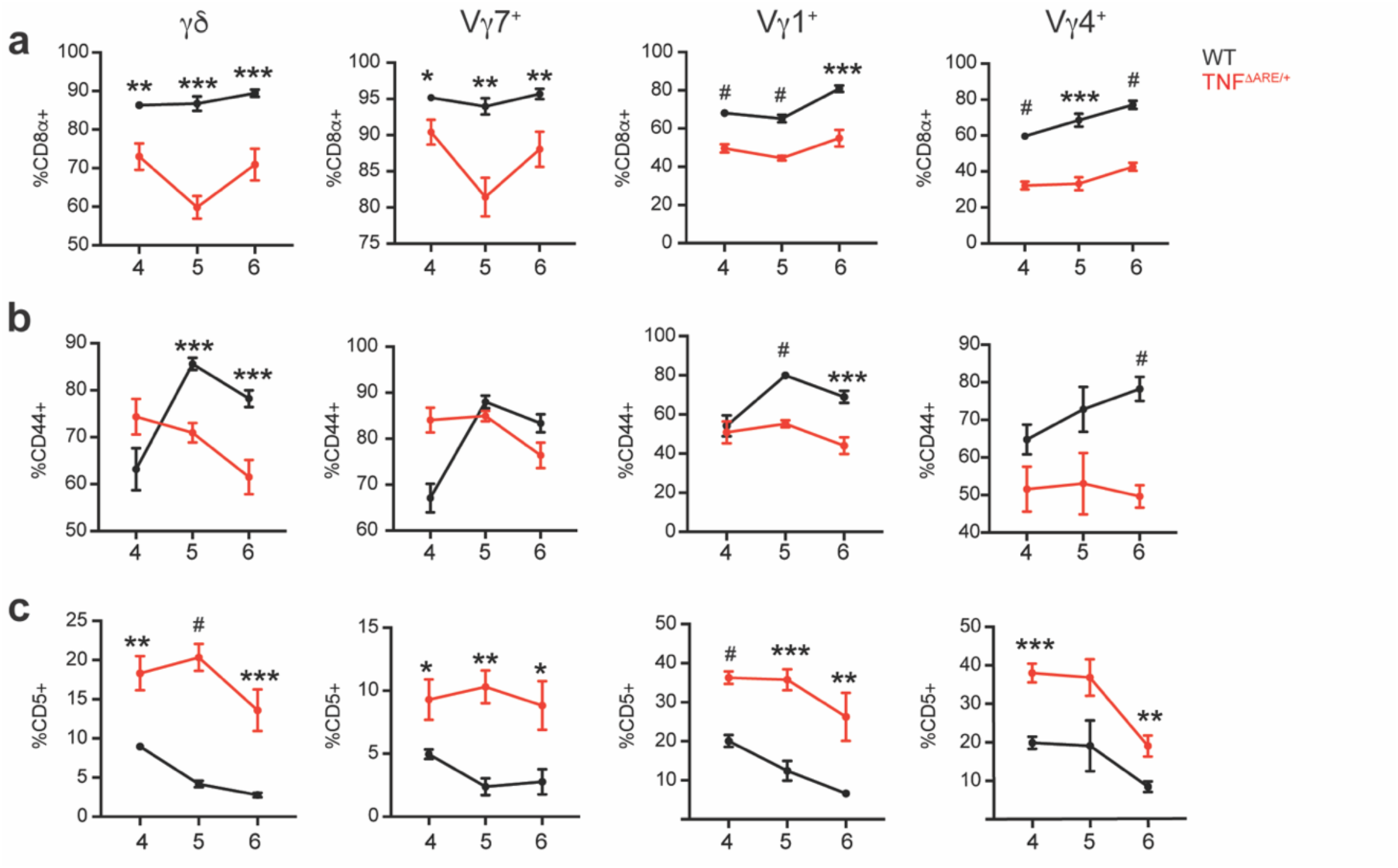
IEL activation and maturation markers are uniformly decreased across Vγ IEL subsets prior to ileitis onset in TNF^ΔARE/+^ mice. Frequency of (a) CD44+, (b) CD8α+, (c) CD5+ in various γδ IEL subsets of WT and TNE^ΔARE/+^ mice from age week 4 to 6. n = 4-11. All data are shown as mean ± SEM from at least 2 independent experiments. Statistical analysis: unpaired t-test. “P<0.05, **P<0.01, ***P< 0.001, #P< 0.0001.

**Supplementary Figure 7:**
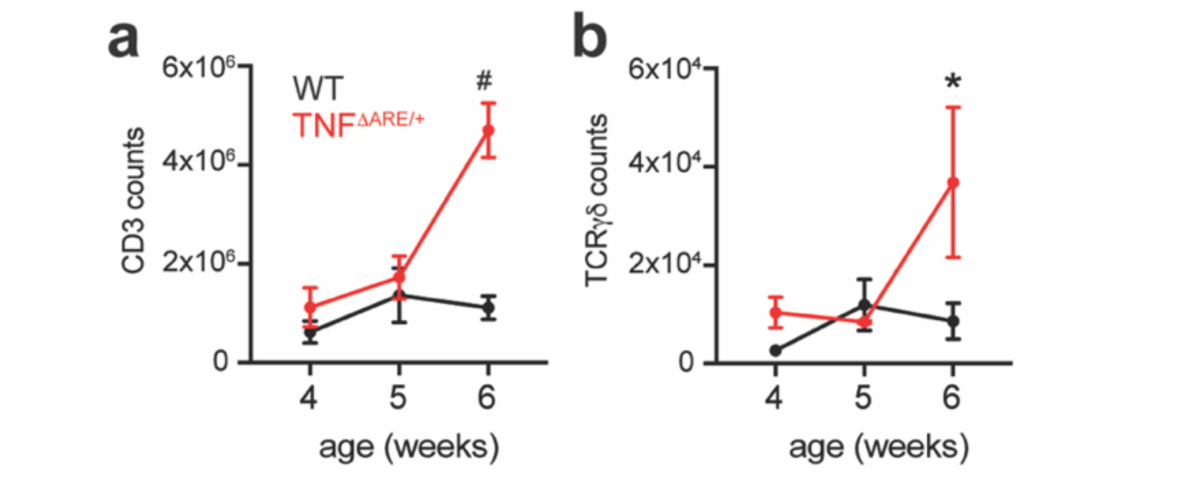
γδ T cell number is increased in the MLN in 6-week-old TNF^ΔARE/+^ mice. (a) Absolute counts of CD3+ and (b) TCRγδ+ in the MLN of WT and TNF^ΔARE/+^ mice from age week 4 to 6. n = 4-9. All data are shown as mean ± SEM from at least 2 independent experiments. Statistical analysis: unpaired t-test.*P<0.05, #P< 0.0001.

**Supplementary Figure 8.**
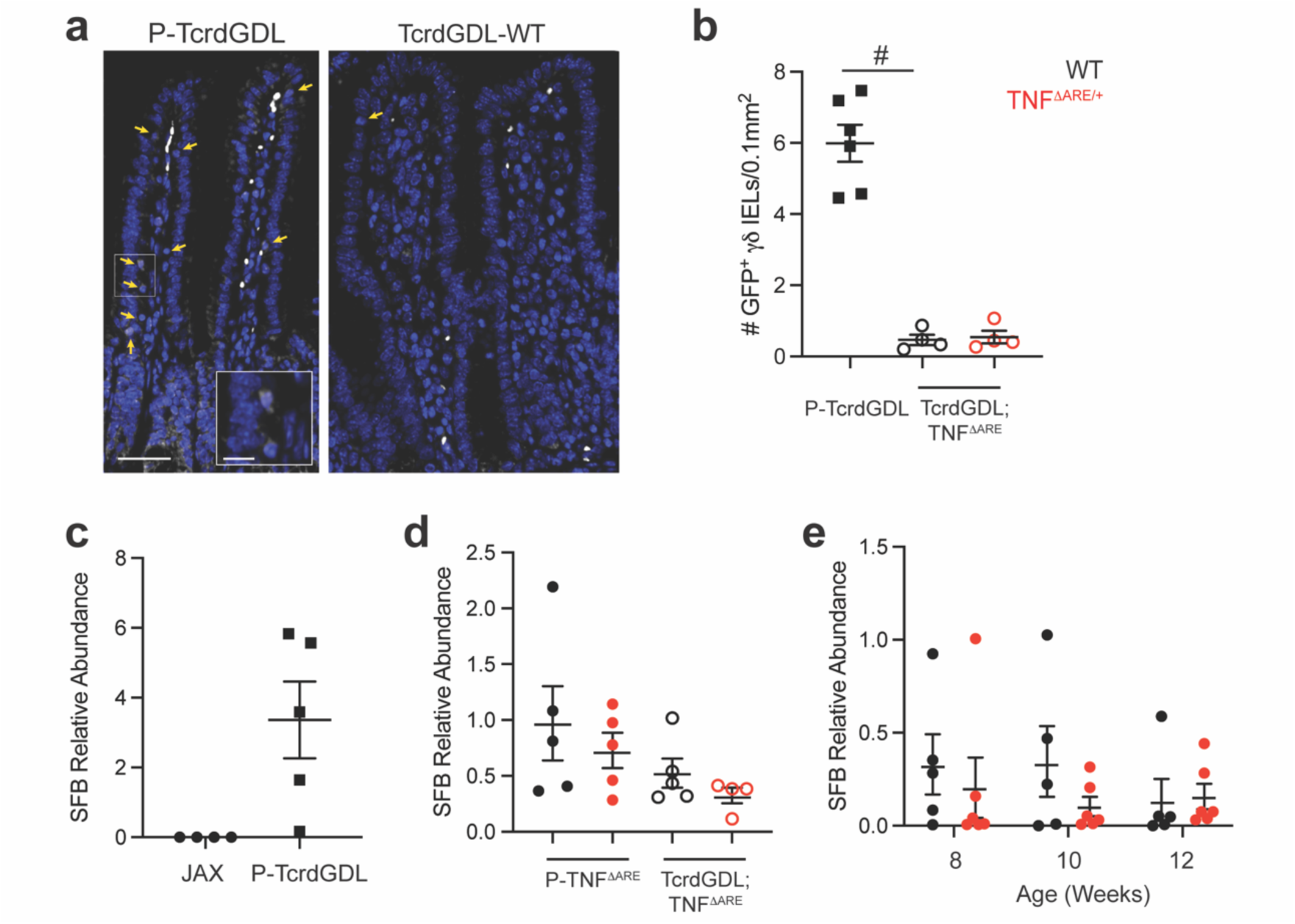
The presence of segmented filamentous bacteria does not correlate with the onset of ileal inflammation in TNF^ΔARE/+^ mice or accelerated disease in TcrdGDL; INF^ΔARE/+^ mice. (a) Immunostaining and (b) morphometric analysis of GFP^+^ γδ IELs (white, yellow arrows) in 6-week-old parental (P)-TcrdGDL mice, TcrdGDL; TNF^ΔARE/+^ and TcrdGDL; TNF^ΔARE/+^ mice. Nuclei are shown in blue. Scale bar = 40 µm, inset = 10 µm. The relative abundance of fecal SFB in (c) 8-12-week-old JAX mice and P-TcrdGDL mice, (d) 6-week-old P-TNF^ΔARE/+^ and TcrdGDL; TNF^ΔARE/+^ mice, and (e) p-TNF^ΔARE/+^ mice during the onset and progression of ileal inflammation. n=4-5. All data are shown as mean ± SEM from at least 2 independent experiments. Each data point represents an individual mouse. Statistical analysis: (b,d) one-way ANOVA with Sidak’s posthoc test, (c) unpaired t-test, (e) two-way ANOVAwith Tukey’s posthoc test. #P<0.0001.

